# Spike–frequency adaptation is modulated by interacting currents in an Hodgkin-Huxley-type model: Role of the Na,K–ATPase

**DOI:** 10.1101/2020.06.12.148437

**Authors:** Renaud Blaise Jolivet, Pierre J. Magistretti

## Abstract

Spike–frequency adaptation is a prominent feature of spiking neurons. Using a Hodgkin–Huxley–type model, we studied adaptation originating from the Na,K–ATPase electrogenic pump and its evolution in presence of a medium–duration calcium–dependent potassium channel. We found that the Na,K–ATPase induces spike–frequency adaptation with a time constant of up to a few seconds and interacts with the calcium–dependent potassium current through the output frequency, yielding a very typical pattern of instantaneous frequencies. Because channels responsible for spike–frequency adaptation can interact with each other, our results suggest that their meaningful time courses and parameters can be difficult to measure experimentally. To circumvent this problem, we developed a simple phenomenological model that captures the interaction between currents and allows the direct evaluation of the underlying biophysical parameters directly from the frequency vs. current curves. Finally, we found that for weak stimulations, the pump induces phasic spiking and linearly converts the stimulus amplitude in a finite number of spikes acting like an inhibitory spike–counter. Our results point to the importance of considering interacting currents involved in spike–frequency adaptation collectively rather than as isolated elements and underscore the importance of sodium as a messenger for long–term signal integration in neurons. Within this context, we propose that the Na,K–ATPase plays an important role and show how to recover relevant biological parameters from adapting channels using simple electrophysiological measurements.

## Introduction

Spike–frequency adaptation is a generic term, which covers all the physiological mechanisms by which the output frequency of a neuron decreases upon continuous presentation of a stimulus. It is exhibited by most neurons and is a widespread phenomenon present in peripheral and central systems of both vertebrates and invertebrates. Spike-frequency adaptation plays a functional role in a variety of phenomena such as forward masking, high-pass filtering and response selectivity [1–7]. From the modeling perspective, it is a necessary mechanism in quantitative neuronal models if we are to capture the firing behavior of cortical neurons and to connect different stimulation regimes [8–12].

Spike–frequency adaptation may originate from several different processes most of which are well–known and having been extensively studied in *in vitro* preparations, as well as in computational models (see e.g. [13, 14]), and most neurons exhibit adaptation at several time scales [11]. Among these, the current generated by the Na,K–ATPase electrogenic pump has relatively recently emerged as a potentially important player with roles in cellular memory formation [15] and synaptic transmission [16], but it is usually ignored in modeling studies because of its absence from the traditional formulation of neuronal biophysics within the formalism established by Hodgkin and Huxley. It is however essential for developing models that go beyond neurons to include glial cells, and their role in brain energy metabolism [17–19]. For these reasons, we undertook the analysis of adaptation that may originate from the activity of this pump in a Hodgkin-Huxley model adapted from [18]. Following sustained activation, sodium accumulates in the neuronal intracellular space and can reach very high concentrations, up to 100 mM in active spines, to be compared with a concentration at rest of ~8–15 mM [20, 21]. High intracellular sodium concentrations increase the activity of the electrogenic Na,K–ATPase pump, which is responsible for an outward current since it extrudes three sodium ions versus two potassium ions at each cycle. In sensory neurons, the Na,K–ATPase induces an adapting current [22–24], but its role in spike-frequency adaptation is not universal. For instance, no primary contribution of the Na,K–ATPase in spike-frequency adaptation was found in hypoglossal motoneurons [25]. The pump also consumes one ATP molecule per cycle, linking the neuronal dynamics to brain energy metabolism and imaging signals [17, 18, 26–28]. We also intended to explore the potential interactions between currents responsible for spike–frequency adaptation. The pattern of instantaneous frequencies typically observed in *in vitro* preparations is the cumulative result of multiple currents [11, 13]. While it is convenient to assume that these currents add up in a relatively simple way, it is not clear if this assumption is actually justified since those currents could interact through the output frequency even if they rely on different biophysical signals or act at different time scales. Interaction of processes over multiple time scales may impact the estimation of relevant parameters and may have important computational consequences [29].

Here, we investigate these two questions using a Hodgkin–Huxley–type model. We show how the electrogenic Na,K–ATPase pump induces spike–frequency adaptation and how it interacts with another adapting channel. A phenomenological model is developed that captures this interaction. We show how the latter model can be used to predict *f–I* curves (frequency vs. current) as well as the relevant time constants. Finally, we provide mechanical insights into how the pump can induce specific discharge patterns like phasic spiking. The model is briefly described in the next section.

## Materials and Methods

### Hodgkin–Huxley–type model

The neuron model is written within the Hodgkin–Huxley framework [30]. The single compartment membrane voltage *V* is given by the balance equation:

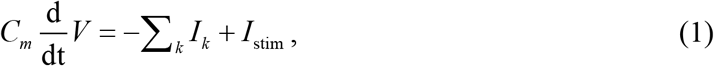

where C_m_=1 μF/cm^2^ is the membrane capacitance and *I*_stim_ is the applied current (in μA/cm^2^). The other currents 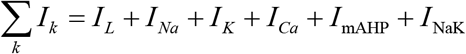 follow the dynamics as introduced by Wang, Heinrich and Schuster [31, 32]. Currents include the typical leak current *I_L_* = *g_L_*(*V* − *E_L_*), the spike generating sodium current 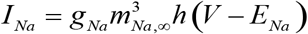 and the delayed–rectifier potassium current *I_K_*=*gKn^4^*(*V-E_K_*) with *g*_*x*_ the conductance of each channel type and *E*_*x*_ the corresponding reversal potential (*x* standing for *L*, *Na* or *K*). The gating variables *m_Na,∞_*, *h* and *n* describe the opening and closing dynamics of voltage–gated sodium and potassium channels. Here, *m_Na,∞_* is assumed to follow the voltage instantaneously:

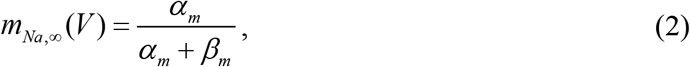

with *α_m_* = −0.1(*V* + 33) /{exp[−0.1(*V* + 33) −1}and *β_m_* = 4 exp[(−*V* + 58)/ 12] [32]. The kinetics of variables *h* and *n* are described by:

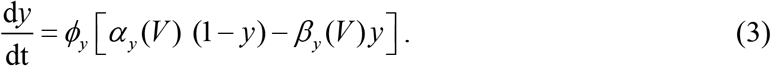

Additional parameters and functions are: *α_h_* = 0.07 exp[− (*V* +50 10)], *β_h_* = (exp [−0.1(*V* + 20)] +1)^−1^, *α_n_* = −0.01(*V* + 34)]/ (exp [−0.1(*V* + 34)] −1), *β_n_* =0.125·exp[−(*V*+44)/25], *ϕ*_*h*_ = *ϕ*_*n*_ = 4, *g*_*Na*_ = 45 mS/cm^2^, *g*_*K*_ = 18 mS/cm^2^, *g*_L_ = 0.1 mS/cm^2^ and *E_K_*=−89mV.

The model also includes a high–threshold calcium current (*I*_Ca_) and a calcium–gated potassium current which, following the literature, we refer to as a mAHP current (*I*_mAHP_) [13]. The calcium current is given by 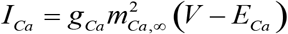 with *m_Ca_*_,∞_ = 1 /{1+ exp [− (*V* + 20) 9]}, *g*_*Ca*_ = 0.005 mS/cm^2^ and *E*_*Ca*_ = 120 mV. The calcium–gated potassium current is given by *I*_mAHP_ = *g*_mAHP_ [Ca^2+^(Ca^2+^ + *K*_*D*_)] (*V* − *E*_*K*_) with *g*_mAHP_ = 5 mS/cm^2^ and *K*_D_=0.03 mmol/L. The internal calcium dynamics is modeled as a leaky–integrator:

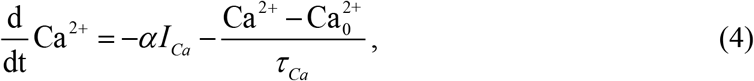

with α=*F^−1^S_m_*/*V_n_*, *F*=9.64853·10^7^ mC/mol, where *F* is the Faraday constant, *S_m_*/*V_n_*=9·10^4^ cm^−1^ the ratio between membrane surface and cell volume [18], 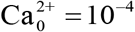mmol/L, and τ_Ca_ = 80 ms the time constant for calcium extrusion.

In addition, the sodium dynamics is described by:

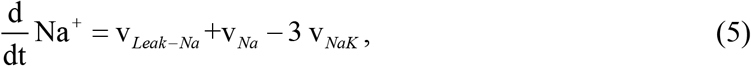

where v_*Na*_=−αI_Na_ describes the sodium movements through voltage-gated channels. The activity of the Na,K–ATPase is given by [18, 31]:

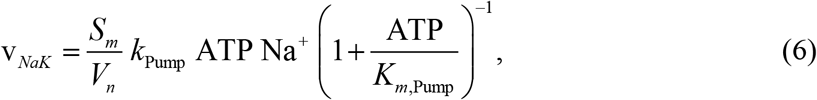

where *k*_Pump_=0.29·10^−9^ cm·L·ms^−1^·mmol^−1^ and *k*_m,Pump_=0.5 mmol/L are constants, and ATP=2.2 mmol/L is the concentration of adenosine triphosphate in the cytoplasm. The leak term is simply given by *v*_Leak-Na_=−α g_Leak-Na_(*V-E_Na_*) with *g*_Leak-Na_=0.0019 mS/cm^2^. To maintain the model at a reasonable level of complexity, the pump activity is measured relatively to the baseline sodium concentration, thus dividing the resulting current in *steady–state* and *dynamic* components [33]. The current generated by the sodium leak channel and the current generated by the pump at baseline sodium concentration are absorbed in the leak current *I_L_*. Therefore, *I*_*NaK*_ is set to zero for baseline intracellular sodium concentration Na_0_^+^=8 mmol/L:

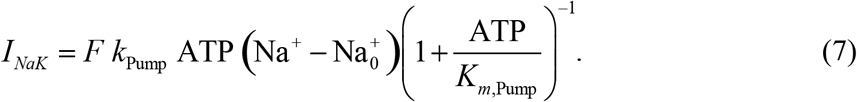

The reversal potential for sodium follows the fluctuations of the intracellular sodium concentration:

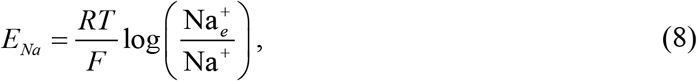

where Na_e_^+^=150 mmol/L is the extracellular sodium concentration and RT/F=26.73 mV. Finally, the reversal potential for the leak current is given by *E*_*L*_ = −78.8 + 0.12 · *E*_*Na*_.

### Phenomenological model

The phenomenological model we use to describe the interaction between the two adapting currents is based on the universal model of spike–frequency adaptation proposed by Benda and Herz [14]. The instantaneous frequency *f* is written:

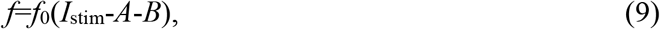

where *f*_0_ is the frequency at stimulus onset as a function of stimulus amplitude. *A* and *B* are adaptation variables representing the two adapting currents. The evolution of *A* and *B* is given by:

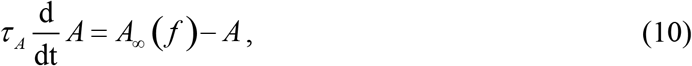

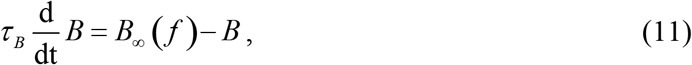

where τ_A_ (respectively τ_B_) is the typical time constant of the adaptation variable *A* (respectively *B*). *A*_∞_ (*f*) and *B*_∞_ (*f*) are the driving forces of the two adaptation variables.

Following Benda and Herz [14], we solve this system at steady–state to find:

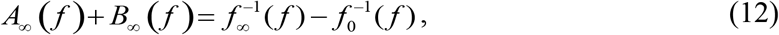

where 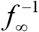 is the inverse of the steady–state frequency as a function of the actual frequency *f*. For the sake of simplicity, we will assume that the two mechanisms are driven by a fraction of the total driving force 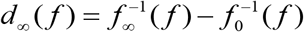 yielding:

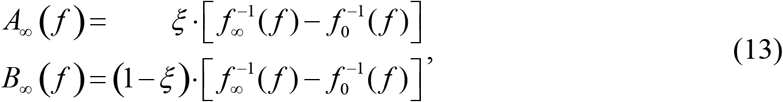

with 0 ≤ ξ ≤ 1 a parameter to be determined. For parameter estimation, the phenomenological model is simply fitted to a single instantaneous frequency curve. Note that this model implicitly assumes temporal averaging [14]. Therefore, it is only valid as long as the actual frequency *f* is above the minimal frequency at which the two mechanisms effectively contribute to spike–frequency adaptation (1/τ_Ca_ = 12.5 Hz in the present case).

## Results

To study the spike–frequency adaptation that may originate from the activity of the electrogenic Na,K–ATPase pump and its interactions with other currents, we simulated the response of an Hodgkin–Huxley–type model to very long sustained tonic stimulation. The model includes an Na,K–ATPase electrogenic pump (*I*_NaK_) plus a calcium–gated potassium current (*I*_mAHP_). Model details can be found in the preceding section.

### The Na,K–ATPase induces spike–frequency adaptation and interacts with the mAHP current

When subjected to long sustained tonic stimulation, the model displays dual time scale spike– frequency adaptation. As the neuron repetitively discharges (Fig. 1A), both intracellular sodium and calcium concentrations rise above baseline level (Fig. 1B). As a consequence, the corresponding currents *I*_NaK_ and *I*_mAHP_ increase, each one inducing spike–frequency adaptation with a different temporal dynamics. At stimulus onset, the instantaneous frequency quickly drops by a significant amount in correspondence with the initial buildup of the calcium concentration and with a typical time constant in the tens of milliseconds range. After this initial phase, the instantaneous frequency slowly decreases to the steady–state frequency with a typical time constant of about ten seconds (Fig. 1C and inset). When the mAHP current is considered in isolation (*k*_Pump_ = 0), it induces spike–frequency adaptation with a typical timescale of a few tens of milliseconds, quasi equivalent to the fast component in the full model (see inset). Removing the effect of the pump does not however completely abolish the slow component since accumulation of sodium in the intracellular space affects the neuronal dynamics through the reversal potential of sodium channels. The resulting slow component is much shorter than when both currents are active indicating that *I*_NaK_ significantly contributes to the late phase of adaptation. Alternatively, the electrogenic pump in isolation induces spike–frequency adaptation with a typical timescale of a few seconds up to about 14s (*g*_mAHP_ = 0; inset). The combination of both currents produces a pattern of instantaneous frequencies very similar to what is typically observed in experimental recordings (see e.g. [34]).

**Figure 1.**
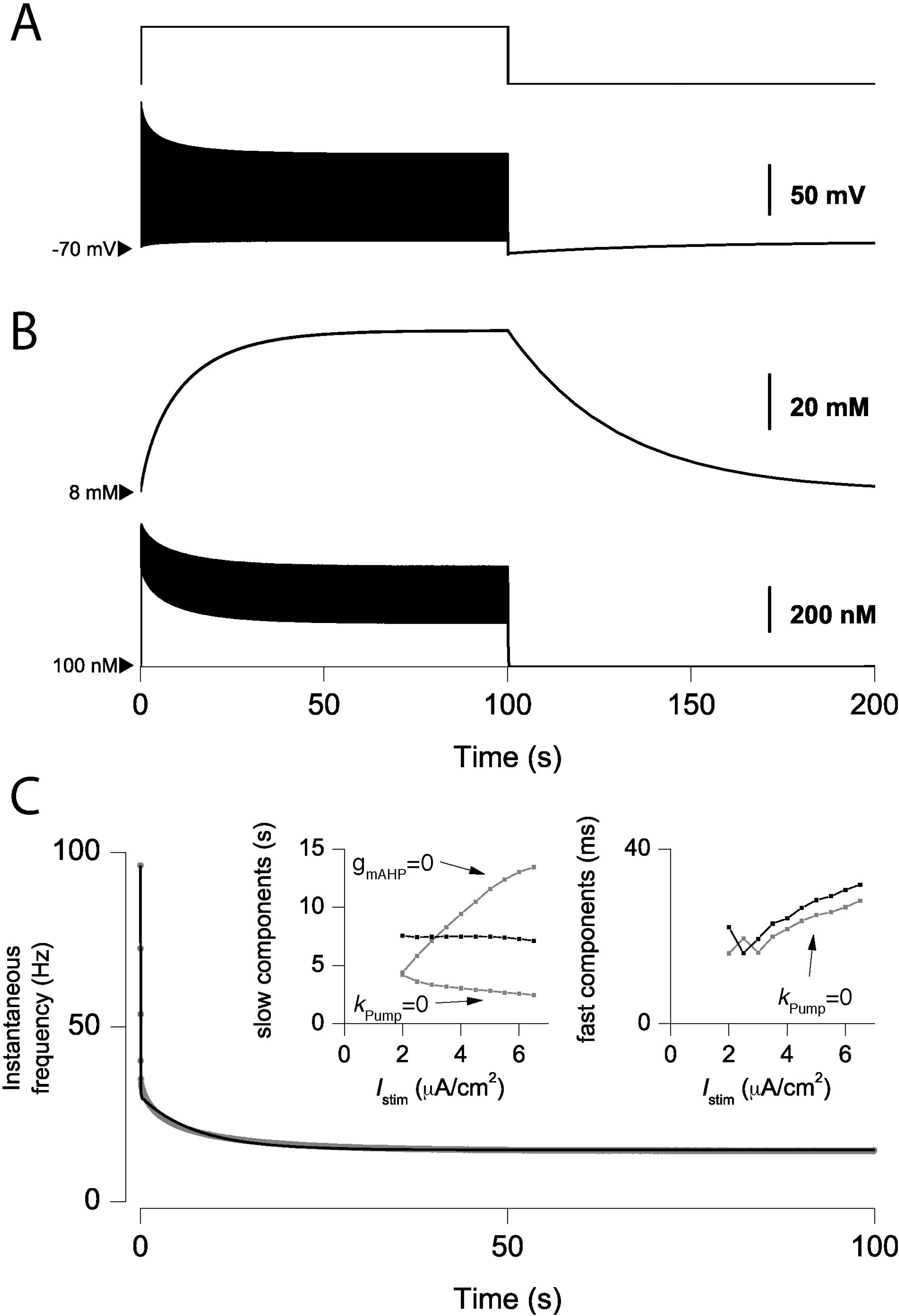
*I*_NaK_ induces spike–frequency adaptation and interacts with *I*_mAHP._ **A.** From top to bottom, applied current (*I*_stim_=2.5 μA/cm^2^) and voltage response. **B.** Intracellular Na^+^ (top) and Ca^2+^ concentrations (bottom). In A and B, arrowheads indicate the baseline concentration/voltage. **C.** Instantaneous frequency during stimulation (symbols) and fitted double exponential (solid line). In inset, slow and fast time constants of the instantaneous frequency are plotted versus the stimulus amplitude *I*_stim_ for the full model (black line), when the calcium–dependent potassium channel is blocked (*g*_mAHP_=0) and when the effect of the electrogenic pump is removed (*k*_Pump_=0). Fast components were omitted when the overall frequency was below the range where calcium accumulates between consecutive spikes (1/τ_Ca_=12.5 Hz).

The accumulation of sodium in the intracellular space also affects the dynamics of the spike mostly by changing the reversal potential for sodium. As sodium accumulates (Fig. 1B), action potentials rise and decay slower and peak lower (Fig. 2A). As a consequence, sodium entry during the action potential is strongly reduced (Fig 2B). On the contrary, the driving force associated with the calcium current is increased and calcium entry is slightly but significantly increased (Fig 2C).

**Figure 2.**
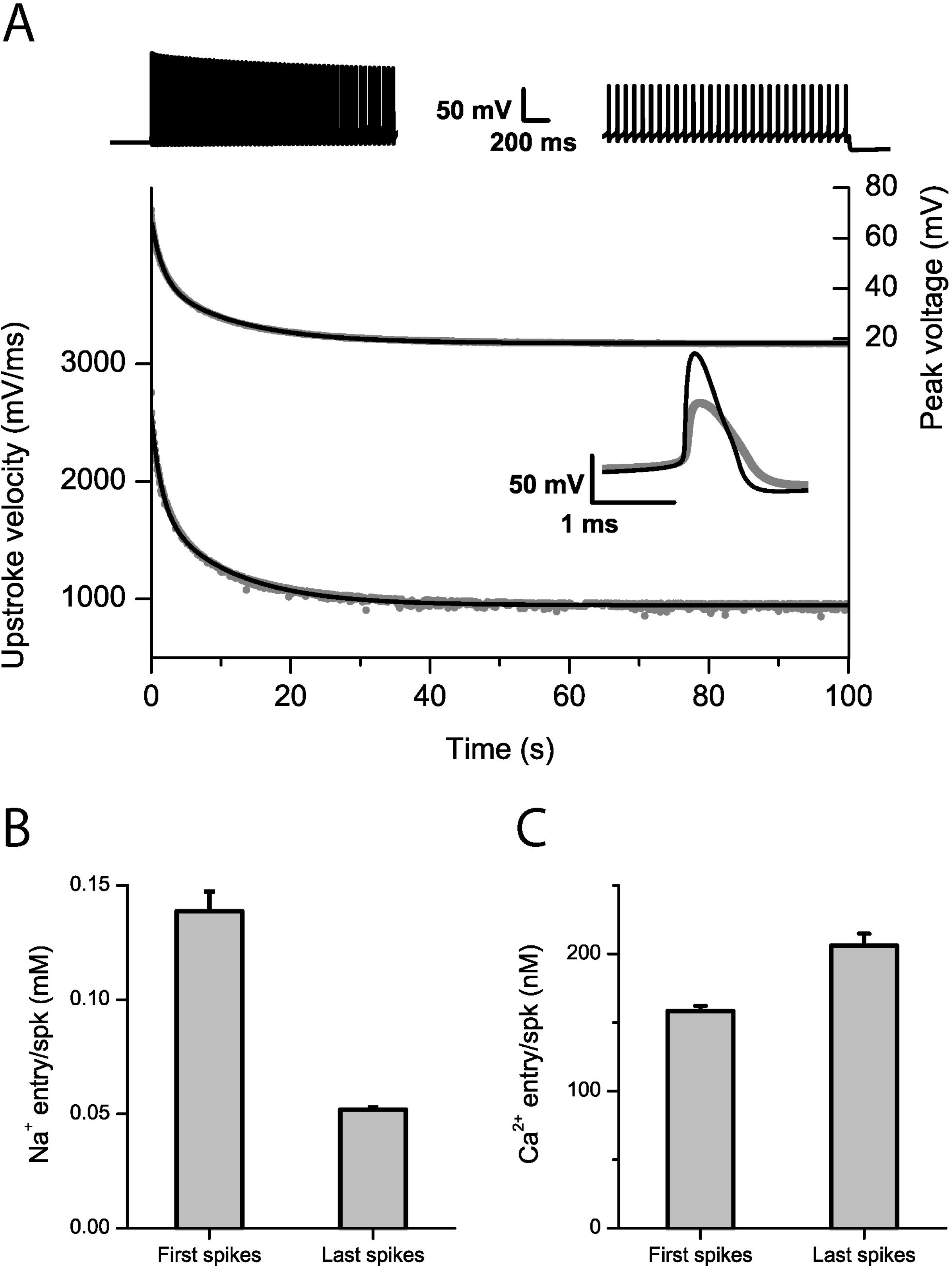
Accumulation of sodium alters the action potential dynamics and sodium entry. **A.** Peak voltage and upstroke velocity during the action potential (symbols) are significantly reduced by the accumulation of sodium in the neuron over the course of a 100 sec long stimulation (*I*_stim_=4 μA/cm^2^). The first few and last spikes are displayed on top while the shape of the first (black) and last (gray) spikes are overlaid in the inset. Both the peak voltage and upstroke velocity decrease can be fitted with a double exponential with similar time constants (black lines; τ_fast_=1.57 sec, τ_slow_=11.2 sec and τ_fast_=1.46 sec, τ_slow_=10.7 sec for the peak voltage and upstroke velocity respectively). **B.** Sodium entry during the action potential is significantly reduced when comparing the first and last ten action potentials (independent t-test; p > 0.05). **C.** On the opposite, calcium entry during the action potential is mildly but significantly increased (independent t-test; p > 0.05).

Following these observations, it would seem natural to conclude that early adaptation can be identified with *I*_mAHP_ while late adaptation can be identified with *I*_NaK_. However, while the fast component is not significantly affected by the presence or absence of the pump (Fig. 1C right inset), the slow component is strongly affected by the pump and does not follow the typical stimulus–dependence displayed when the pump alone contributes to adaptation. When *I*_mAHP_ is blocked, the effective adaptation time constant increases with increasing current like for several other mechanisms including *I*_mAHP_ [14, 32]. On the contrary, it is roughly constant when *I*_NaK_ and *I*_mAHP_ are both present. This suggests that an interaction is taking place between the two currents during the late phase of adaptation as also illustrated by the biphasic calcium dynamics in Fig. 1B. This can be explained as follows. While *I*_NaK_ and *I*_mAHP_ do rely on different biophysical signals, respectively sodium and calcium concentrations, both depend on spiking and in turn, affect spiking. Hence, the two currents interact through the instantaneous frequency, and this interaction affects the neuronal dynamics in the late phase of adaptation after sodium has increased significantly compared to its basal level. As a direct consequence, blocking the fast channel affects the late time course of the slow channel and could thus bias estimates of its temporal dynamics in relation to physiological conditions.

### Phenomenological model of interacting channels

In order to capture the interaction between adapting channels and to circumvent the related parameter estimation issue, we developed a phenomenological model based on the universal model of spike–frequency adaptation proposed by Benda and Herz [3, 14]. Our instance of the model is detailed in *Materials and Methods*. Briefly, the instantaneous frequency *f* is written:

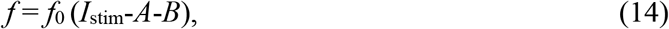

where *f*_0_ is the frequency at stimulus onset as a function of stimulus amplitude *I*_stim_. *A* and *B* are adaptation variables representing the two adapting currents *I_mAHP_* and *I_NaK·_* Both *A* and *B* evolve as leaky–integrators with time constants τ_A_ (respectively τ_B_) and are driven by a fraction *ξ* (respectively 1− ξ) of the horizontal difference *d*_∞_ measured between steady–state frequency f∞ and onset frequency *f*_0_ (Fig. 3A). This formalism allows a direct estimation of meaningful parameters that would not be possible by using solely the instantaneous frequency curve [14, 32]. In particular, τ_A_ and *τ*_*B*_ can be identified with the time constants of the underlying biophysical mechanisms.

**Figure 3.**
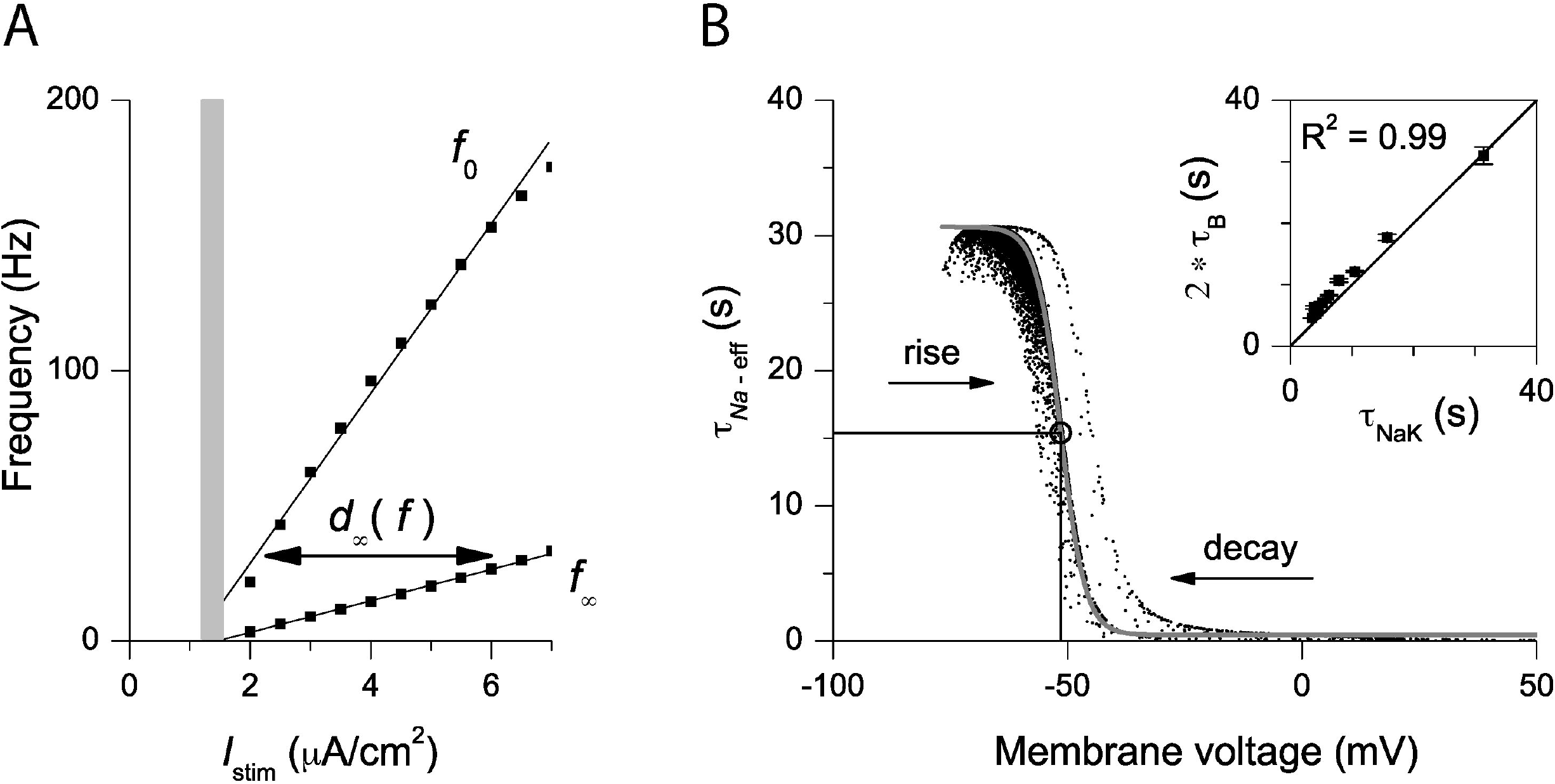
Phenomenological model of spike–frequency adaptation. **A.** Onset and steady–state frequencies (*f*_0_ and *f*_∞_). At frequency *f*, abstract variables *A* and *B* are driven by a fraction of the horizontal distance *d*_∞_ between *f*_0_ and *f*_∞_. The grey area denotes the region where phasic spiking occurs. **B.** Effective sodium time constant τ_Na-eff_ ranges between 9.7 ms and 30.7 sec. Data corresponding to interspike intervals and action potential rising phases are fitted by a Boltzmann function τ_Na-eff_=τ_min_+(τ_max_-τ_min_)/(1+exp[*V*-*V*_0_]/δV) with parameters τ_min_=440 ms, τ_max_=30.6 sec, *V*_0_=−51.5 mV and δV=2.6 mV (grey line). The effective time constant is centered around the threshold for spike initiation (≈V_0_) and time constant (τ_min_+τ_max_)/2=15.5 sec (circle). During the action potential decaying phase, the effective time constant is larger than during the rising phase because voltage–gated sodium channels are closed. The effective time constant τ_Na-eff_ is evaluated after linearizing the sodium reversal potential appearing in *v*_Leak-*Na*_+*v*_Na_ and reorganizing the terms in Eq. (5). Inset shows a comparison between the true time constant for sodium extrusion τ_Na__K_ and the time constant estimated using the proposed method.

To evaluate parameters τ_A_, τ_B_ and *ξ*, the output frequency *f* of the abstract model is simply fitted to a sample instantaneous frequency curve after evaluation of *f*_0_ and *f*_∞_. In the case of our Hodgkin–Huxley model, this procedure persistently yields τ_A_=80±9 ms, τ_B_=15.5±0.7 sec and ξ=0.43±0.02 (mean ± sem). Now, τ_A_ can be compared with the time constant for calcium extrusion set in the Hodgkin–Huxley model τ_Ca_=80 ms and we note the excellent match between the phenomenological model prediction τ_A_ and the set value τ_Ca_. For τ_B_, the situation is slightly more complicated since contributions of leak and voltage–gated sodium channels make the sodium time constant fluctuate between the time constant for sodium extrusion by the Na,K-ATPase and the much faster time constant of activated voltage–gated channels. Hence, τ_B_ cannot be readily compared to the sodium extrusion time constant (τ_NaK_=31.4 sec) but should rather be compared to the *effective* time constant. An estimate of this effective time constant including all the contributions shows that its dependence on voltage follows a Boltzmann function centered at threshold for spike initiation (−51.5 mV) and around an effective value τ_Na-eff_=15.7 sec (Fig. 3B; see Materials and Methods for how we evaluate τ_Na-eff_). Again, we note the good match between τ_B_ and τ_Na-eff_. Since the abstract model identifies the effective time constant for sodium dynamics and since the latter is about half the value of τ_NaK_ (see legend of Fig. 3), we can retrieve the value of τ_NaK_ by simply doubling the value that we obtain for τ_B_. This simple method yields good predictions for the time constant of the Na,K-ATPase (Fig. 3B inset). This approximation is valid as long as τ_NaK_ is much larger than the time constants playing a role in fast sodium transients. This is usually the case with τ_NaK_ observed to be in the range of seconds to minutes [20, 21].

The phenomenological model also allows the prediction of instantaneous frequency curves (Fig. 4A). Predictions are excellent except at low frequencies, where it is expected to fail, i.e. when the frequency drops below the value at which calcium–gated potassium channels do contribute to adaptation (1/τ_Ca_=12.5 Hz; see *Materials and Methods*). Finally, the phenomenological variables *A* and *B* retain the qualitative behavior of their Hodgkin–Huxley counterparts. The interaction between channels and the temporal dynamics of relevant concentrations are captured in their dynamics as illustrated in Fig. 4B (compare the dynamics of *A* and *B* with the dynamics of sodium and calcium in Fig. 1B).

**Figure 4.**
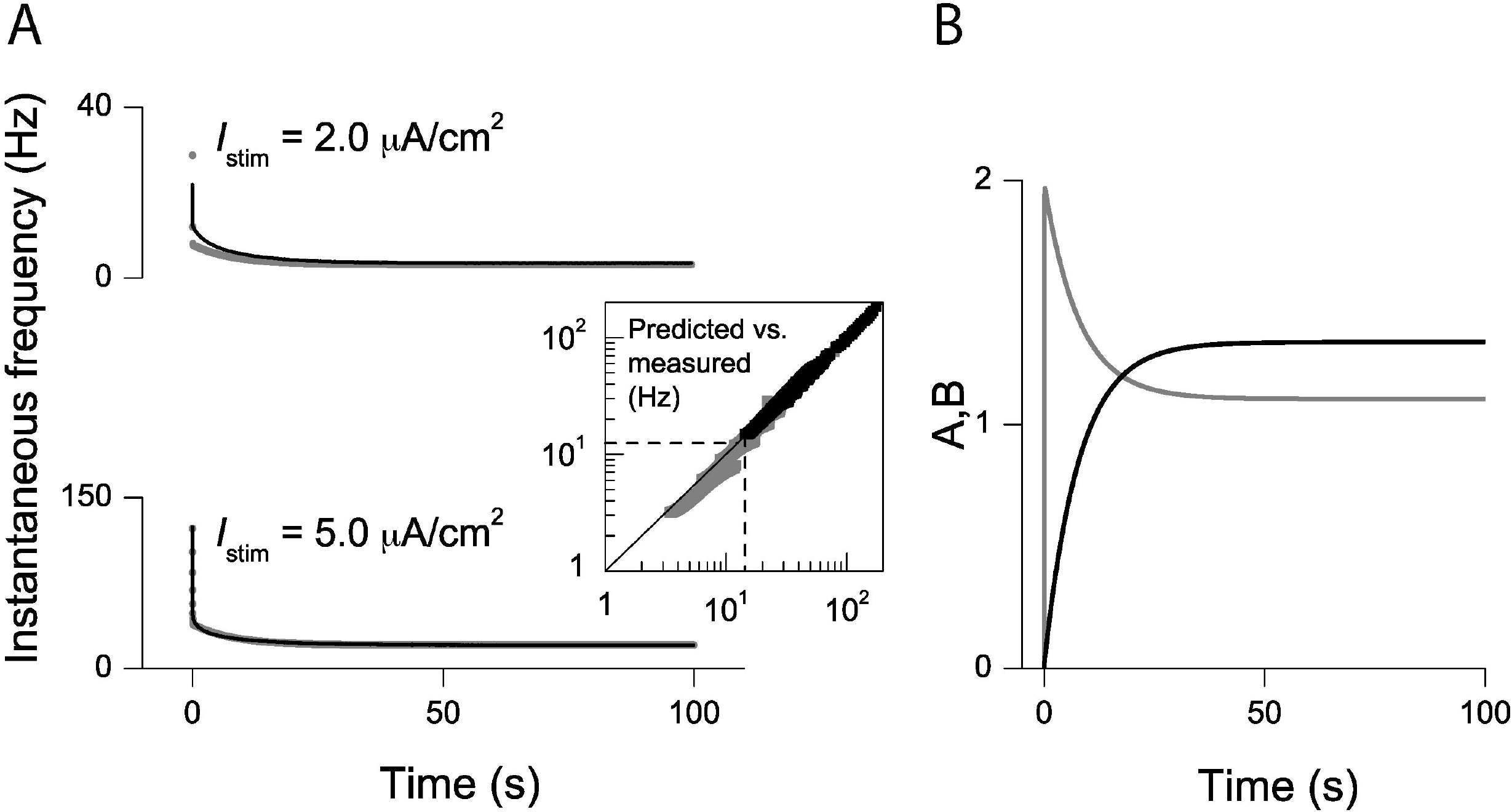
Phenomenological model dynamics. **A.** Prediction of *f–I* curves using the abstract model (black line) are compared to data (symbols) for two different values of the stimulating current. Predictions are excellent above the frequency at which calcium accumulates between consecutive spikes (bottom graph; 1/τ_Ca_=12.5 Hz signaled by a dotted line in inset; black symbols in inset; R^2^>0.99), but relatively poor below this frequency (top graph; grey symbols in inset; R^2^>0.96). **B.** Time course of adaptation variables *A* (grey) and *B* (black) during typical stimulation (*I*_stim_=4μA/cm^2^).

### The Na,K–ATPase induces phasic spiking

For sufficiently strong stimulation intensities (*I*_stim_>1.55 μA/cm^2^), the model displays a continuous *f–I* curve (Fig. 3A). However, the model is not readily a type I neuron, i.e. with a continuous *f–I* curve from zero frequency on [35]. There exists a narrow regime of stimulation just above threshold where *I*_NaK_ induces phasic spiking (1.2 ≤ *I*_stim_ ≤ 1.55 μA/cm^2^; Fig. 3A). Spiking stops after a few emitted spikes even though the stimulation is maintained (Fig. 5A). This type of behavior cannot be obtained with the mAHP current alone. Indeed, while calcium entry through high–threshold channels is entirely dependent on spiking, sodium continues to flow in the neuron through voltage–gated channels even after spiking has stopped and the system eventually reaches a regime where the current generated by the pump exactly balances the stimulating current (Fig. 5B). For stronger stimulations (*I*_stim_>1.55 μA/cm^2^), stimulation plus strong activation of voltage–gated sodium channels (*I*_stim_*+I_Na_*) override the inhibition due to *I*_NaK_ and the model behaves roughly as a type I neuron. In this phasic spiking regime, stimulus amplitude is linearly converted in a finite number of spikes and the pump acts like an inhibitory “spike–counter” (Fig. 5C).

**Figure 5.**
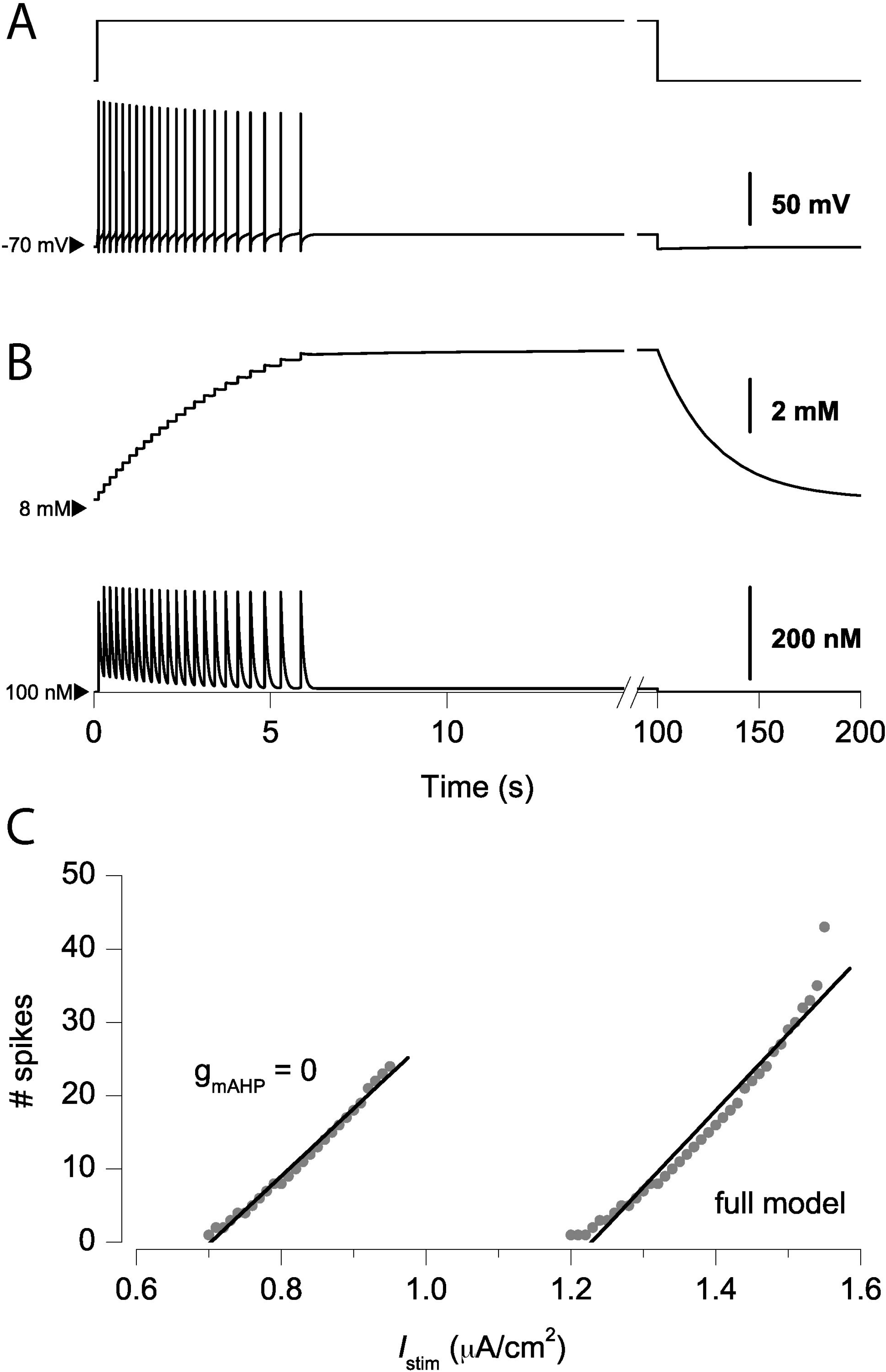
*I*_NaK_ induces phasic spiking. **A.** From top to bottom, applied current (*I*_stim_=1.45 μA/cm^2^) and voltage response. **B.** Intracellular Na^+^ (top) and Ca^2+^ concentrations (bottom). In A and B, arrowheads indicate the baseline concentration/voltage. **C.** The number of spikes produced in the phasic spiking regime is plotted versus the applied current *I*_stim_ for the complete model and when *I*_mAHP_ is blocked (symbols). In both cases, the relation between the stimulus amplitude and the number of spikes produced is approximately linear (solid lines; in both cases R^2^>0.95 and slope=98.5 spk·(μA/cm^2^)^−1^).

## Discussion

Using Hodgkin–Huxley formalism, we investigated the role of the Na,K–ATPase electrogenic pump on neuronal dynamics and more specifically, in spike–frequency adaptation in presence of a calcium–dependent potassium current, another channel inducing adaptation with a much shorter typical time constant. We have demonstrated here how currents inducing spike– frequency adaptation can collectively interact with each other through the output frequency, even when they act at very different time scales (Fig. 1). This interaction is not restricted to the specific choice of currents we made in this report but is rather a generic mechanism. It should take place between any number of currents relying directly or indirectly on spikes for activation, like M–type currents, ATP–dependent potassium currents, electrogenic pumps or sodium and calcium–activated potassium currents. Interestingly, interacting processes have been proposed as an efficient way to model power-law adaptation, a form of adaptation that may have far-reaching computational consequences [29]. A consequence of this interaction is that pharmacologically blocking a channel will affect the time course of the remaining currents, therefore possibly biasing measures of their temporal dynamics. To circumvent this problem, we developed a simple phenomenological model of spike–frequency adaptation based on earlier work by Benda and Herz [14]. The latter model captures the essence of this interaction. It allows the evaluation of the parameters of underlying mechanisms using only onset and steady–state *f*–*I* curves and extends the results of Benda and Herz to two interacting channels (Fig. 3 and 4). This simple formalism can be used to measure typical extrusion time constants of multiple channels contributing to spike–frequency adaptation without prior knowledge of specific neuronal dynamics. There are no *a priori* reasons to think that this approach would fail for three or more interacting channels. The interaction between channels also seems to produce input-independent effective adaptation time constants (Fig. 1), a potentially interesting behavior that will require further study.

We also found that the Na,K–ATPase induces phasic spiking in a limited range of stimulation conditions (Fig. 5). Spiking stops after a few emitted spikes even though the stimulation is maintained and the pump acts like an inhibitory spike–counter, linearly converting stimulus amplitude in a finite number of spikes. This Na,K–ATPase–dependent behavior had been previously reported by Sokolove and Cooke in crayfish tonic stretch receptor neurons [22] and is similar to the reported role of the Na,K-ATPase as an integrator of spike number in rhythmically active *Drosophila* neurons [15]. This effect is specific to the sodium dynamics, involving a subtle balance between early subthreshold activation, high density of voltage–gated sodium channels and the activity of the Na,K-ATPase. A parallel can be drawn with another sodium–specific dynamics where subthreshold activation of voltage–gated sodium channels is important, the so–called EPSP amplification [36, 37]. Our analysis provides mechanistic insights into the role played by the Na,K–ATPase in burst termination and burst–length control in brain stem motoneurons [33] and midbrain dopaminergic neurons [38]. Regulation of the Na,K–ATPase could then have important computational consequences in dopaminergic pathways [38].

Several processes have been neglected in our approach. We can safely ignore fast afterhyperpolarizing currents since they only induce adaptation within a few spikes after stimulus onset [13]. Sodium–calcium exchangers were neglected as well but because of the difference between intracellular sodium and calcium concentrations, their impact on our results should be negligible. Adaptation due to slow afterhyperpolarizing calcium-dependent channels is taking place on a much longer time scale. However, the exact origin of this current is still speculative [39]. There is also evidence that sodium-dependent potassium channels and slowly inactivating sodium channels can contribute to the late phase of adaptation [34, 40]. In light of our results, it could well be that no single channel is responsible for the current observed in the late phases of adaptation, but rather that a group of channels acting cooperatively lead to this current. The impact of sodium–channels inactivation is left for future study, as it would largely complicate the present analysis.

Our results are not fundamentally dependent on exact parameter values. The principal parameter for which different values are equally likely is the pump constant *k*_Pump_. We chose its value so as to obtain an overall time constant for sodium extrusion in agreement with experimental observations [21]. Nevertheless, we varied *k*_Pump_ over a realistic range and, while quantitative results do depend on its exact value, we found no significant qualitative differences with the results presented here. In particular, phasic spiking was observed for a large variety of parameters sets.

Finally, our results underscore the importance of sodium as a messenger for long–term signal integration [41]. They suggest a role potentially more important than usually considered for the Na,K–ATPase in signal processing and notably because of the slow time constant for sodium extrusion, at low frequencies accessible *in vivo*. Our findings are consistent with observations in the peripheral nervous system and subcortical structures [33, 38]. Whether the Na,K–ATPase significantly affects the dynamics of cortical neurons in a similar way remains to be determined. One may however hypothesize that it is of importance in thin dendrites where sodium concentration can rise up to very high values [21] and where, concordantly, metabolic demand following activation is very important [18, 28, 42, 43].

## Acknowledgments

This work was supported by grants from the Swiss National Science Foundation (31003A_170079), the European Commission (H2020 FET-Open IN-FET 862882) and the Australian Research Council (GA2481) to RBJ. The authors wish to thank Dr. Igor Allaman for advice.

## Notes

### Competing Interest Statement

The authors have declared no competing interest.

